# The structure of the OmpA/Pal protein of *Anaplasma phagocytophilum*

**DOI:** 10.1101/2023.06.15.545016

**Authors:** I.T. Cadby

## Abstract

Peptidoglycan associated lipoprotein (Pal) and Outer Membrane Protein A (OmpA), interact with the outer membrane and peptidoglycan in Gram-negative bacteria, conferring structural integrity to the bacterial cell and functioning in cell division. Both OmpA and Pal proteins have moonlighting roles as virulence factors, facilitating infection and host-pathogen interactions in a range of bacteria. The OmpA-like protein of *Anaplasma phagocytophilum*, a tick-borne pathogen that infects a wide range of hosts, seems to function primarily as a virulence factor, since this bacterium lacks a peptidoglycan cell wall. Here we present crystal structures of the OmpA-like protein of *A. phagocytophilum*, demonstrating that this protein has amino acid insertions that confer flexibility. This insertion is also found in the OmpA-like proteins of other pathogens, related to *A. phagocytophilum*. Whether this flexibility is reflective of any adaptations for host-pathogen interactions remains to be determined but, since the OmpA-like proteins of *Anaplasma* species are current targets for vaccine development, might have importance for these efforts.

## INTRODUCTION

*Anaplasma* is a genus of Gram-negative bacteria that lead obligate intracellular lifestyles (being entirely reliant on eukaryotic host cells for their survival and replication) and which infect ticks and a wide range of vertebrates including wild and domestic animals, livestock, and humans ^1^. Generally recognised as infecting the blood cells of their vertebrate hosts, different species of *Anaplasma* have varied host species and host cell tropisms. For example, *Anaplasma marginale (Am)* preferentially infects the erythrocytes of ruminants whereas *Anaplasma phagocytophilum (Ap)*, a zoonotic agent with a wide vertebrate host range, shows preference for neutrophils. *Anaplasma* infection can cause disease within their vertebrate hosts and tick-borne fever and bovine anaplasmosis cause significant losses to livestock industries across the globe^2,3^. Infection of humans with *Ap* causes human granulocytic anaplasmosis: an emerging and occasionally fatal disease^4^.

Peptidoglycan associated lipoprotein (Pal) and outer membrane protein A (OmpA) are conserved in most Gram-negative bacteria^5,6^. These proteins are integrated into or bound to the outer membrane and bind to peptidoglycan (PG), tethering these major structural components to one another. OmpA comprises two main protein domains: a beta barrel outer membrane protein domain (OMP), that is inserted into the outer membrane, and a soluble OmpA-like domain, that residues in the periplasm and binds to PG. OmpA anchorage between the outer membrane and PG is regarded as having an important role in conferring structural stability to the bacterial cell and integrity to the periplasmic space, but has other functions including porin activity, facilitating the movement of solutes^7^. Pal comprises an OmpA-like domain (also referred to as PAL), and, as a bacterial lipoprotein, has an N-terminal signal peptide lipobox that enables attachment to the outer membrane via acylation. Pal has roles in cell division and contributes to invagination of the cell membrane and septum development^8 - 10^. Both OmpA and Pal have moonlighting roles as virulence factors in some bacteria^11,12^. OmpA and Pal proteins are implicated in adhesion, biofilm formation, and immune modulation; and have been identified as vaccine candidates, in a range of bacteria.

The genomes of *Ap*, and some other Rickettsiales (the bacterial Order to which *Ap* belongs), lack the full complement of genes necessary for PG biosynthesis^13-15^. Indeed, fluorescent labelling of PG structures in various Rickettsiales, supports that *Ap* does not synthesise PG^16^. However, other Rickettsiales genomes, such as *Am* and *Orientia tsutsugamushi*, encode complete or partially complete PG biosynthesis pathways and are able to synthesise PG, with some members producing an diminutive PG cell wall^14,16^. Despite lacking PG, *Ap* retains vestiges of the PG pathway and also encodes PG binding proteins, including the OmpA-like protein of *Ap* (OmpA^*Ap*^, APH_0338). In line with the loss of PG, OmpA^*Ap*^ functions as an invasin to facilitate infection of mammalian host cells^17-19^. Intriguingly, all evidence demonstrates that OmpA^*Ap*^ recognises sialic acid and fucose moieties of mammalian cell surface glycoproteins, and can bind the tetrasaccharide sialyl-Lexis-X. Consistent with this, the OmpA of *Am* also has glycan binding activity linked to adhesin and invasion activities^20^. Both *Ap* and *Am* express highly variable complements of outer membrane proteins, process called antigenic variation, that confers immunoprotection to these pathogens^21-24^. Considering the importance of OmpA of *Ap* and *Am* as virulence factors, and that these proteins are encoded by single genes in these respective bacteria (and therefore do not undergo antigenic variation), they are regarded as promising vaccine and therapeutic targets to combat disease^25^.

To gain insights into how *Anaplasma* OmpA-like proteins are adapted to function as virulence factors and glean information that could aid the development of interventions targeting these proteins, we solved the crystal structure of OmpA^*Ap*^ to a resolution of 1.9 Å and assessed its potential to bind sialyl-Lewis-X and other sugars. Our results demonstrate that OmpA^*Ap*^ is most similar to Pal and contains amino acid insertions, generating an extended loop region proximal to its PG binding site. This insertion also occurs in *Am* and some other Rickettsiales. Whilst *in vitro* binding of recombinant OmpA^*Ap*^ to sialyl-Lewis-X could not be detected, these results present a basis for further investigation and highlight that structural flexibility for *Anaplasma* OmpA proteins might be a consideration in vaccine development.

## METHODS

### Cloning, protein expression and purification

The APH_0338 gene, corresponding to amino acids 10-205 of OmpA^*Ap*^, was amplified from *Ap* strain HZ gDNA using Q5 DNA polymerase (New England Biolabs). Cloning into pET-28a and a modified pET41 was performed using restriction free PCR-based methods^26^. The cloning yielded two T7 inducible overexpression vectors: pET_APH0338N, encoding APH_0338 lacking its N-terminal signal peptide and with an N-terminal 6 x histidine tag and pET_APH0338C, encoding the same gene sequence but with a C-terminal 8 x histidine tag. The constructs were transformed into *E. coli* DE3 and then used for protein over-expression. Briefly, 10-20 ml cultures of *Escherichia coli* strains were grown in LB broth at 37 °C overnight and used as the inoculum for large-scale (1-2 L) cultures in auto-induction medium. Cultures were grown at 37 °C with shaking for approximately 3 hours until an OD_600_ reading of 0.6-0.8 had been reached, at which point the temperature was reduced to 18 °C. After an additional 18 hours of growth the cultures were harvested by centrifugation and cell pellets were then frozen for later use in protein purification.

To purify OmpA^*Ap*^ proteins, cell pellets (10-25 g) were resuspended in Buffer A (20 mM imidazole-HCl pH 8.0; 50 mM Tris-HCl pH 8.0; 400 mM NaCl; 2 mM BME) supplemented with 10 ug/ml hen egg lysozyme and 0.02 % Tween 20 and lysed by sonication. Lysates were clarified by centrifugation at 20,000 rpm and then passed over nickel agarose beads in gravity flow columns. Beads were washed extensively with Buffer A, then with 15 column volumes of Buffer A supplemented with 10 % Buffer B (as Buffer A but with 400 mM imidazole, pH 8.0), then with 15 column volumes of Buffer A supplemented with 20 % Buffer B, before final washes with Buffer B. The eluates were analysed by SDS-PAGE, indicating that high purity OmpA was collected in the Buffer B washes. Proteins were dialysed into Buffer C (Tris-HCl, pH 8.0; 250 mM NaCl; 1 mM BME) and then further purified by gel filtration in Buffer C using a Superdex 200 26/60 column. OmpA^*Ap*^ proteins typically eluted at a volume consistent with a protein with a MW of 20 kDa. This monodisperse peak was pooled and concentrated to 20 mg/ml with a 10 kDa MWCO ultrafiltration device (Sartorius).

### Protein crystallography

Crystallization of OmpA^*Ap*^ proteins was achieved by mixing equal volumes of concentrated purified protein and various Molecular Dimensions crystallization screens in sitting drop trays and then incubation at 18 °C. Crystals of C-terminally tagged OmpA^*Ap*^ grew after four weeks in condition A10 of screen JCSG. These small protein crystals were briefly transferred into a drop of mother liquor supplemented with 25 % ethylene glycol prior to snap freezing in liquid nitrogen. X-ray diffraction data was collected at the Diamond Light Source Synchrotron and processed using the CCP4 and Phenix software suites. The structure of OmpA^*Ap*^ was initially solved using by molecular replacement using the structure of Pal from *Burkholderia cepacian* (PDB:5LKW) as the search query. This model was refined using Phenix, manual adjustments using Coot software, and the PDB-redo server. Attempts were also made to crystallize OmpA in the presence of sialyl Lewis X, fucose, and sialic acid. Trials involved protein supplemented with a 2-fold molar excess of the proposed ligand but yielded no crystals. Attempts were also made to soak these ligands into apo-OmpA^*Ap*^ crystals: mother liquor was supplemented with 2 mM of the ligand and added to OmpA^*Ap*^ protein crystal containing drops. Crystals were harvested at various time points, from 5 minutes after addition up until 24 hours after addition before harvesting as described above. None of these strategies yielded electron density maps consistent with ligand bound protein.

## RESULTS

### Architecture of *A. phagocytophilum* OmpA

The OmpA^*Ap*^ protein is comprised of an N-terminal signal peptide and lipobox, a central OmpA-like/PAL domain, and a disordered C-terminal stretch of approximately 40 amino acids **(Figure 1A)**. Comparison of the predicted domains of OmpA^*Ap*^ with the known domain architecture of typical OmpA and Pal proteins, such as those of *E. coli* (OmpA^*Ec*^ and Pal^*Ec*^), indicates that OmpA^*Ap*^ lacks a beta-barrel OMP domain. In terms of protein domain architecture, OmpA^*Ap*^ closely resembles Pal^*Ec*^.

**Figure 1:**
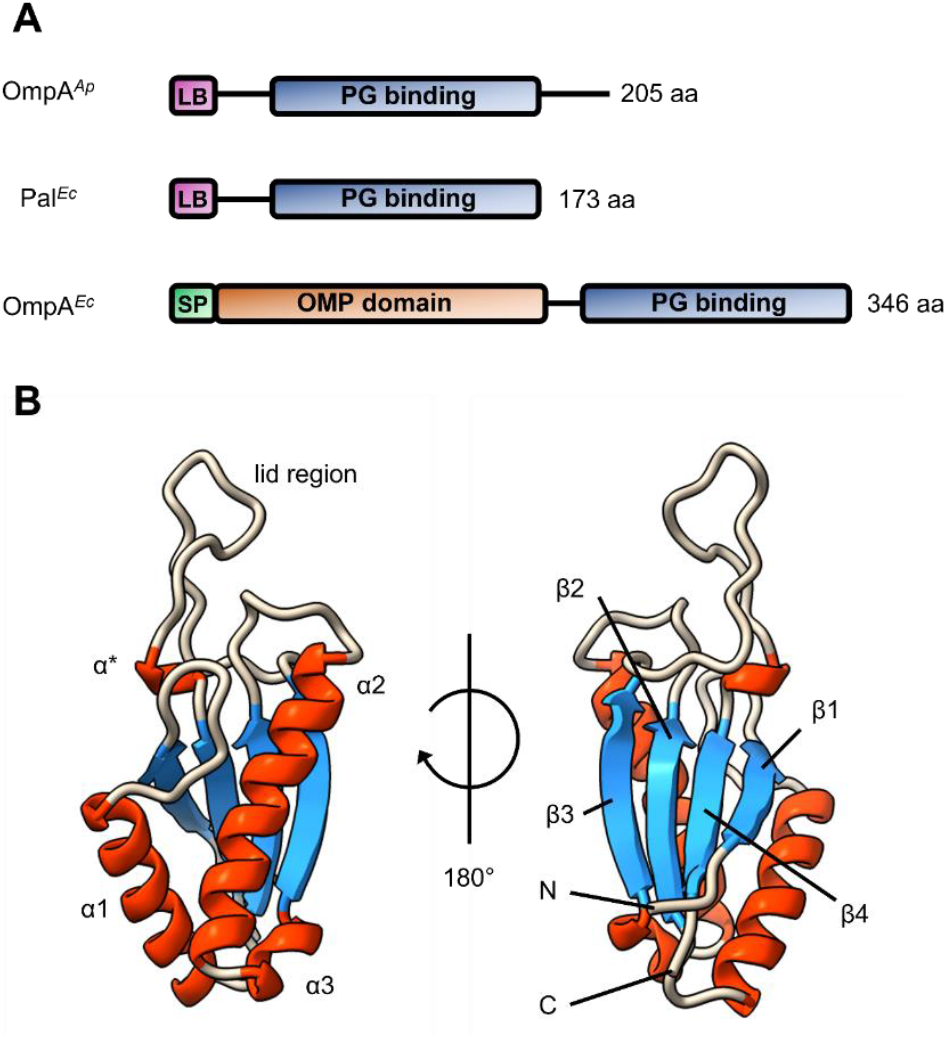
Structural characterisation of OmpA^Ap^. A. Schematic representation of the protein architecture of OmpA and Pal proteins from *E. coli* and *Anaplasma phagocytophilum*. Peptidoglycan binding (OmpA-like/PAL) domains (PG binding), Lipoboxes (LB),Signal peptides (SP), and Outer Membrane Protein (OMP) domains are labelled. B. Cartoon representation of the crystal structure of OmpA^Ap^, solved to 1.9 Å. α* marks a variable length helix.

### Overall structure of Omp^*Ap*^

Crystals of Omp^*Ap*^ were grown by sitting drop vapour diffusion and took approximately 4 weeks to appear. X-ray diffraction data was collected on these crystals, which belong to the orthorhombic C222 space group, and the Omp^*Ap*^ structure was solved by molecular replacement and refined to a resolution of 1.9 Å. Data processing and refinement statistics are displayed in **Table 1**. Two monomers of Omp^*Ap*^ are present in the symmetric unit. The overall fold of Omp^*Ap*^ is similar to that of other OmpA-like/PAL domains and is comprised of an alpha-beta sandwich, ordered as β1-α1-β2-α2-β3-α*-β4, where α* is a variable length helix, discussed below (**Figure 1B)**. Typical of the OmpA-like/PAL fold, adjacent strands β1, β4, and β2 are antiparallel to each other, with β3 being parallel to β2 and completing a β-sheet which associates with α1 and α2. No density was observed for amino acids 1-32 and 165-205, possibly indicating that these regions are disordered, in line with secondary structure predictions. The two monomers pack together with Cys115 from each chain being close enough to form a disulfide (had reducing agent been omitted from protein preparations) and two intermolecular salt bridges formed by Arg75 and Asp116 from each monomer. However, gel filtration experiments with Omp^*Ap*^ suggest the protein is monomeric in solution and so there is no evidence that these interactions have physiological relevance, beyond their role as crystal contacts.

**Table 1:** Data collection and refinement statistics

|  | **OmpA<sup>Ap</sup>** |
| --- | --- |
| **Data collection** |  |
| Spacegroup | C 2 2 2 |
| Cell dimensions |  |
| a,b,c (Å) | 133.4 138.5 34.7 |
| $\alpha, \beta, \gamma$ (°) | 90 90 90 |
| Resolution | 69.3 – 1.90 |
| I/σI | 12.11 2.45) |
| Completeness (%) | 99.65 (97.71) |
| Redundancy | 2.0 (2.0) |
| **Refinement** |  |
| Resolution (Å) | 1.9 |
| No. of reflections | 26011 |
| R<sub>work</sub>/R<sub>free</sub> | 0.1797 / 0.2175 |
| No. of atoms |  |
| Protein | 1908 |
| Ligand/ion | 6 |
| Water | 340 |
| B factors |  |
| Protein | 21.54 |
| Ligand/ion | 15.54 |
| Water | 33.29 |
| RMSD |  |
| Bond lengths (Å) | 0.007 |
| Bond angles (°) | 1.15 |

Superimposing the two monomers of Omp^*Ap*^ in the asymmetric unit onto each other gives an RMSD for equivalent atoms of 0.291 Å for residues 43-164. The two monomers are structurally identical with the notable exception of amino acids 135-150. These residues bridge β3-β4 and form a region we have termed α* or the ‘lid’ (Figure 1B).

### Omp^*Ap*^ retains residues for PG binding

In modelling the Omp^*Ap*^ structure, strong positive *Fo–Fc* density was observed in a pocket formed by coils between β1-α1, β2-α2 and residues from the adjacent beta-sheet and α2. The crystallization conditions that yielded Omp^*Ap*^ crystals contained sodium formate. Accordingly, formate was could be unambiguously modelled into this density (**Figure 2A, B)**. Formate is present in the pockets of both monomers and is co-ordinated by hydrogen bonds with the side chains of Arg104, and the main chain amines of Asp89 and Gly55.

**Figure 2:**
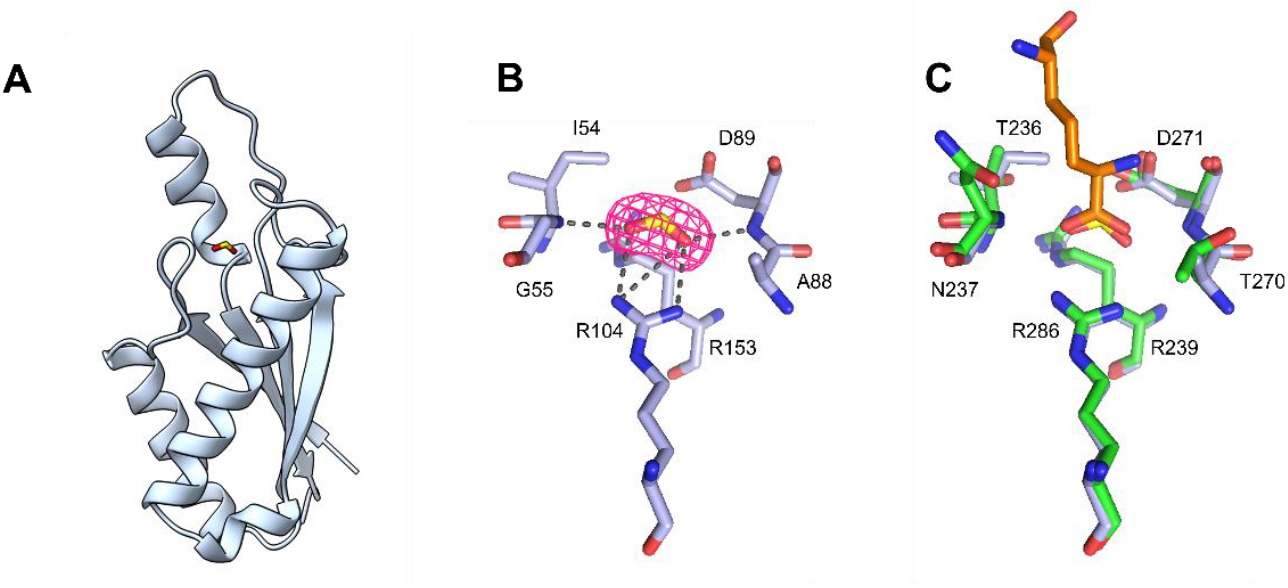
The coordination of formate in the OmpA^*Ap*^ pocket resembles PG binding by other OmpA-like/PAL domains. A. Cartoon representation of OmpA^*Ap*^ with formate modelled into the pocket. B. Fo − Fc omit map demonstrating the presence of density (pink mesh, contoured at 3σ) that corresponds to formate (yellow). Multiple hydrogen bonds (dashed lines) with the peptide backbone (shown as sticks) co-ordinate the ligand. C. Superposition of the OmpA^*Ap*^ - formate complex with an OmpA-like domain from *Acinetobacter baumannii* in complex with a peptidoglycan fragment (PDB:3TD5). The diaminopimelic acid group of peptidoglycan is shown in orange, the OmpA-like domain of *A. baumannii* is coloured green, and OmpA^*Ap*^ in light blue.

Since OmpA-like/PAL domains bind to PG via an equivalent pocket structure we superimposed the structure of Omp^*Ap*^ with that of the OmpA-like domain of *Acinetobacter baumannii* (OmpA^*Ab*^) in complex with a peptidoglycan fragment (**Figure 2C**). In superimposing these two structures the formate that occupies the pocket of Omp^*Ap*^ superimposes onto and mimics the carboxyl group of diaminopimelic acid, a component of the PG stem peptide, that occupies the equivalent pocket of OmpA^*Ab*^. Co-ordination of the carboxyl group of diaminopimelic acid in the OmpA^*Ab*^ pocket is mediated by hydrogen bonds from an Arg side-chain and main chain nitrogen groups that are equivalent to those which co-ordinate formate in the Omp^*Ap*^ structure. Additional hydrogen bonding to diaminopimelic acid is mediated by an Asp residue which also has an equivalent in the Omp^*Ap*^ structure. The comparison of Omp^*Ap*^ with an PG bound OmpA-like/PAL protein indicates that this protein has conserved residues in its pocket that are required for binding to PG. This conservation of residues indicate that Omp^*Ap*^ most likely retains its ancestral PG binding abilities, despite the lack of this macromolecule in *Ap*.

### An extended loop of Omp^*Ap*^ forms multiple conformations

Comparisons between the two Omp^*Ap*^ monomers reveals that the lid adopts two conformational states. In one conformation (chain A) residues 142-153 adopt an α-helical conformation (**Figure 3A)**. However, in the alternative conformation (chain B), this helix shortens, corresponding to residues 149-153, and with residues 142-147 instead adopting a coil conformation. The sum of this restructuring results in the lid region twisting and moving by 14.94 Ä, relative to the Cα of residue 144. These two alternative conformations of Omp^*Ap*^ are each stabilized by crystal contacts so it is not possible to determine how readily the lid forms these two different states in solution. Superposition of Pal from *E. coli* (Pal^*Ec*^, PDB: 1OAP) with these structures indicates that the lid region of Omp^*Ap*^ is extended compared to canonical members of this protein family (**Figure 3A)**. A multiple alignment of Omp^*Ap*^ with the Pal family proteins of other Rickettsiales and representative Gram-negative bacteria demonstrates that this extension is due to the insertion of amino acids in OmpA^*Ap*^ **(Figure 3B)**. This insertion is seen in other Rickettsiales OmpA-like/PAL proteins but varies in length. For example; in *Ap, Am, Ehrlichia chaffeensis*, and *Wolbachia* species, this insertion is six amino acids in length; whereas in *O. tsutsugamushi* it is two amino acids; and selected *Rickettsia* species do not have any amino acid insertion to the lid region of their Pal proteins.

**Figure 3:**
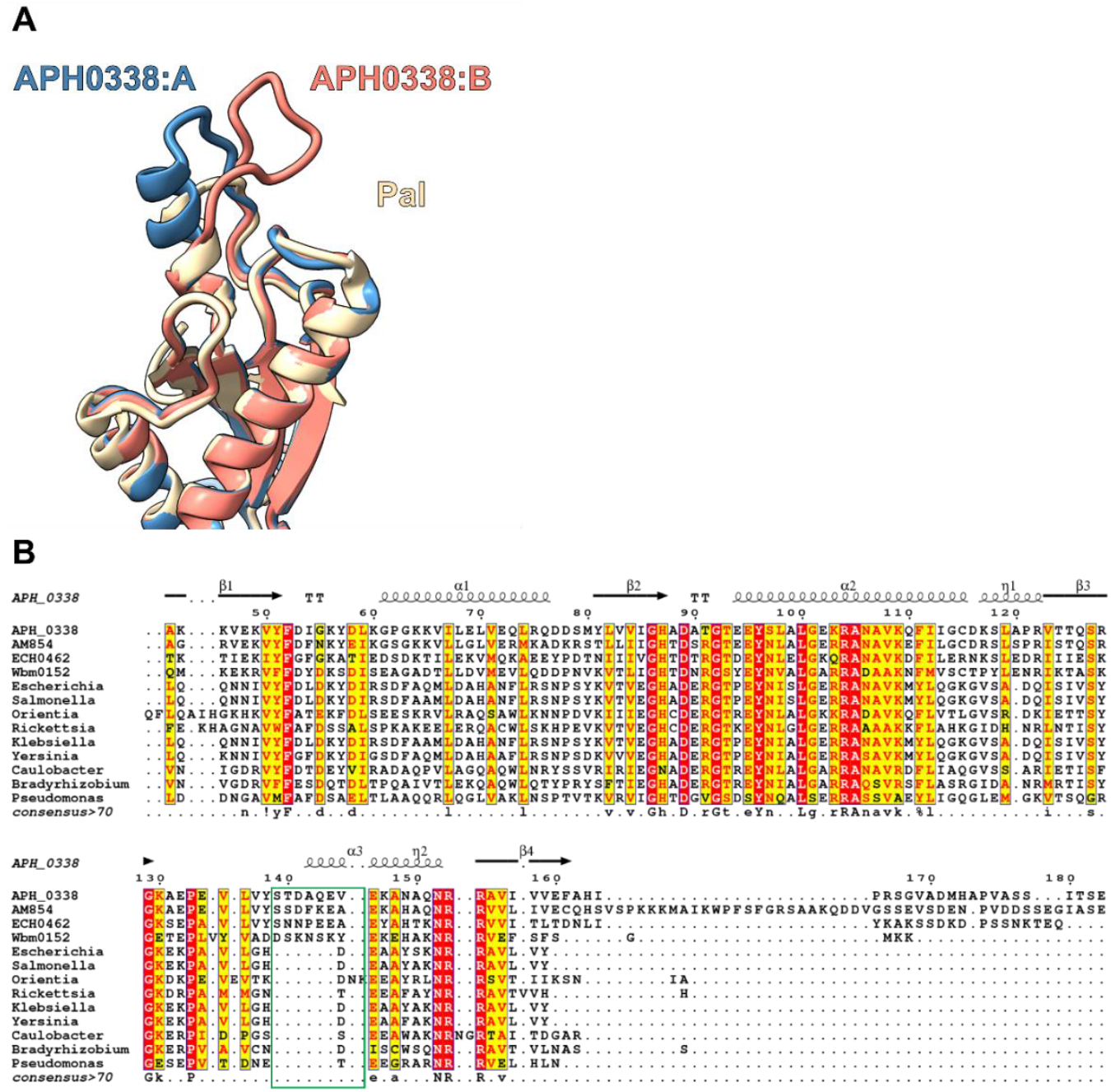
Structure and sequence comparisons between Omp^Ap^ and representative OmpA-like/PAL family proteins of diverse Gram-negative bacteria. A. Superposition of two conformations of Omp^Ap^ and Pal^Ec^. The lid region of these Pal family proteins is shown, detailing the amino acid insertion and flexibility of Omp^Ap^. B. Multiple sequence alignment of OmpA-like/PAL family proteins. The inserted amino acids of the lid region are marked with a green box.

### An extended loop of Omp^*Ap*^

## DISCUSSION

The Omp^*Ap*^ structures presented here demonstrate that the OmpA-like/PAL family proteins of *Ap, Am* and other Rickettsiales contain inserted sequences and are divergent from typical members of this protein family. This insertion provides structural flexibility to OmpA^*Ap*^, proximal to its PG binding site, which is conserved despite the lack of PG in this bacterium. It is unknown whether this switching of conformations occurs in solution, but seems unlikely to impact the potential for Omp^*Ap*^ to bind its ancestral ligand, PG, since similar insertions are seen in OmpA-like/PAL proteins found in Rickettsiales that do synthesise a PG cell wall, such as *Am*. Crucially, this flexibility could influence the ability for antibodies to recognise Rickettsiales OmpA-like/PAL family proteins.

Attempts to measure binding of Omp^*Ap*^ to its host targets, such as sialyl-Lewis-X, by various approaches such as microscale thermophoresis or isothermal calorimetry were unsuccessful, as were attempts to co-crystallize Omp^*Ap*^ with this sugar (data not shown). This is consistent with the proposed hypothesis that Omp^*Ap*^ binds to host surfaces co-operatively with other *Ap* adhesins and/or invasins^18^. The potential role that insertions in OmpA-like/PAL proteins might play in mediating host-pathogen interactions, a conserved function for the OmpA-like/PAL proteins of *Anaplasma*, is unknown.

## Notes

### Competing Interest Statement

The authors have declared no competing interest.

### Summary of Updates

Additional figure added.

